# The 2-oxoglutarate/malate carrier extends the family of mitochondrial carriers capable of FA-activated proton transport

**DOI:** 10.1101/2023.12.14.571682

**Authors:** Kristina Žuna, Tatyana Tyschuk, Taraneh Beikbaghban, Felix Sternberg, Jürgen Kreiter, Elena E. Pohl

**Author notes:** Ludwig Boltzmann Institute for Traumatology, The Research Centre in Cooperation with AUVA, 1200 Vienna, Austria. Institute of Molecular and Cellular Physiology, Stanford University School of Medicine, Stanford, CA 94305, USA.

## Abstract

Metabolic reprogramming in cancer cells has been linked to the mitochondrial dysfunction. Recent studies have suggested the mitochondrial 2-oxoglutarate/malate carrier (OGC) as a potential target for preventing cancer progression. Although OGC is known to be a part of the malate/aspartate shuttle, its exact role in cancer metabolism remains unclear. In this study, we investigated the contribution of recombinant murine OGC to the proton transport by measuring the conductance (G_m_) of planar lipid bilayer membranes reconstituted with OGC. Our results show that OGC significantly increases G_m_ only in the presence of free fatty acids (FAs) and 2,4-dinitrophenol, demonstrating for the first time its involvement in proton transport. We found that (i) the increase in OGC activity directly correlates with the increase in the number of unsaturated bonds of FAs, and (ii) OGC substrates and inhibitors compete with FAs for the same binding site. In addition, we have identified R90 as a crucial amino acid of the binding site for FAs, ATP, 2-oxoglutarate, and malate, which is a first step towards understanding the OGC-mediated proton transport mechanism. Elucidating the contribution of OGC to the uncoupling will be crucial in the design of targeted drugs for the treatment of cancer and other metabolic diseases.

## Introduction

Cancer cells exhibit altered metabolism, relying more on aerobic glycolysis than oxidative phosphorylation for energy production, as well as increased glutaminolysis and oxidative stress ^1,2^. In particular, glutamine-derived α-ketoglutarate (AKG) is converted to citrate by reductive carboxylation, deviating from the usual TCA cycle. The oxoglutarate-malate carrier (OGC) assists by transporting AKG and malate across the mitochondrial membrane ^2^. Recent evidence shows that knockdown of the 2-oxoglutarate/malate carrier (OGC) results in a 75% reduction in ATP production in non-small cell lung cancer ^3^. In addition, a heterozygous OGC knockout reduced the growth of spontaneous lung cancer in mice by 50%, while inhibition of OGC by N-phenylmaleimide resulted in a 50% reduction in melanoma formation in human xenograft models ^4^. The inhibitory effect on the malignant growth has been attributed to the involvement of OGC in the replenishment of NADH required for ATP production via the malate-aspartate shuttle. Neutralization of excessive amounts of reactive oxygen species (ROS) by increasing proton transport across the membrane may be an alternative strategy and has been shown to induce apoptosis in mature tumors or prevent damage early in cancer development and after radiation therapy ^5^. Application of chemical protonophores such as BAM15, niclosamide, or oxyclozanide to *in vitro* and *in vivo* systems resulted in a significant reduction in tumor proliferation^6–8^.

OGC, a member of the mitochondrial solute carrier 25 (SLC25) superfamily, is located in the inner mitochondrial membrane (IMM), where it transports 2-oxoglutarate (α-ketoglutarate) for L-malate or other C4 metabolites ^9^. Notably, the double knockout of OGC is embryonically lethal ^3^. OGC was previously used as a negative control for uncoupling ^10,11^ until Yu et al. observed a significant decrease in mitochondrial membrane potential (MMP) after its overexpression in HEK293 cells ^12^. Later, glutathione transport by OGC was proposed to play a role in reducing of oxidative stress in neuronal cells ^13^, but this function remained controversial ^14^. Since oxidative stress is reduced by mitochondrial uncoupling ^15,16^, the real reason may be OGC-mediated H^+^ transport, which has never been confirmed and further investigated in a well-defined system.

Since the homologs of OGC - adenine nucleotide transporter 1 (ANT1, 28% homology to OGC) and uncoupling proteins 1 (UCP1, 33% homology to OGC) - enhance H^+^ transport in the presence of free fatty acids (FAs) and 2,4-dinitrophenol (DNP) ^17,18^, we hypothesized that also OGC dissipates the IMM proton gradient under similar conditions. Therefore, in this work, we used a well-defined system of planar lipid bilayer membranes reconstituted with OGC to (i) investigate its contribution to FA- and DNP-mediated proton transport, (ii) evaluate the efficiency of OGC inhibitors in reducing the proton transport rate, and (iii) identify amino acids critical for the interaction of OGC with FAs.

Unravelling the physiological function and mechanism of OGC-mediated mitochondrial uncoupling has the potential to provide insight into its involvement in cancer metabolism and aid in the development of targeted drugs for other metabolic diseases.

### Results

### 1. OGC is present in a wide range of murine tissues and cancer cell lines

To understand the physiological role of OGC, we first investigated its tissue expression in mice. Based on immunohistochemical staining experiments and testing the mRNA abundance, OGC has been reported to be ubiquitously expressed in the human body ^19^, and in various rat tissues ^20^. However, mRNA levels are not reliable predictors of protein presence ^21^ and there are no experimental data on OGC protein expression in mice. The main problem in testing the expression of SLC25 superfamily proteins is the lack of specific antibodies capable of distinguishing close homologs. To address this issue, we first validated the anti-OGC (anti-SLC25A11) antibody using inclusion bodies (IBs) of mouse OGC (mOGC) and other SLC25 family members, as well as heterozygous mouse knockout tissues. As shown in Figure S1B, the antibody was specifically bound only to OGC IBs. We found that OGC was expressed on the protein level in all tested murine tissues (Fig. 1A), with an expected decrease in heterozygous knockout samples. OGC was also present at the mRNA level in all tissues (Fig. 1B), and in both cases, the highest expression was found in the heart, brain, and kidney. In addition, we have shown that OGC is expressed at different levels in murine and human cancer cell lines (Fig. 1C).

**Figure 1.**
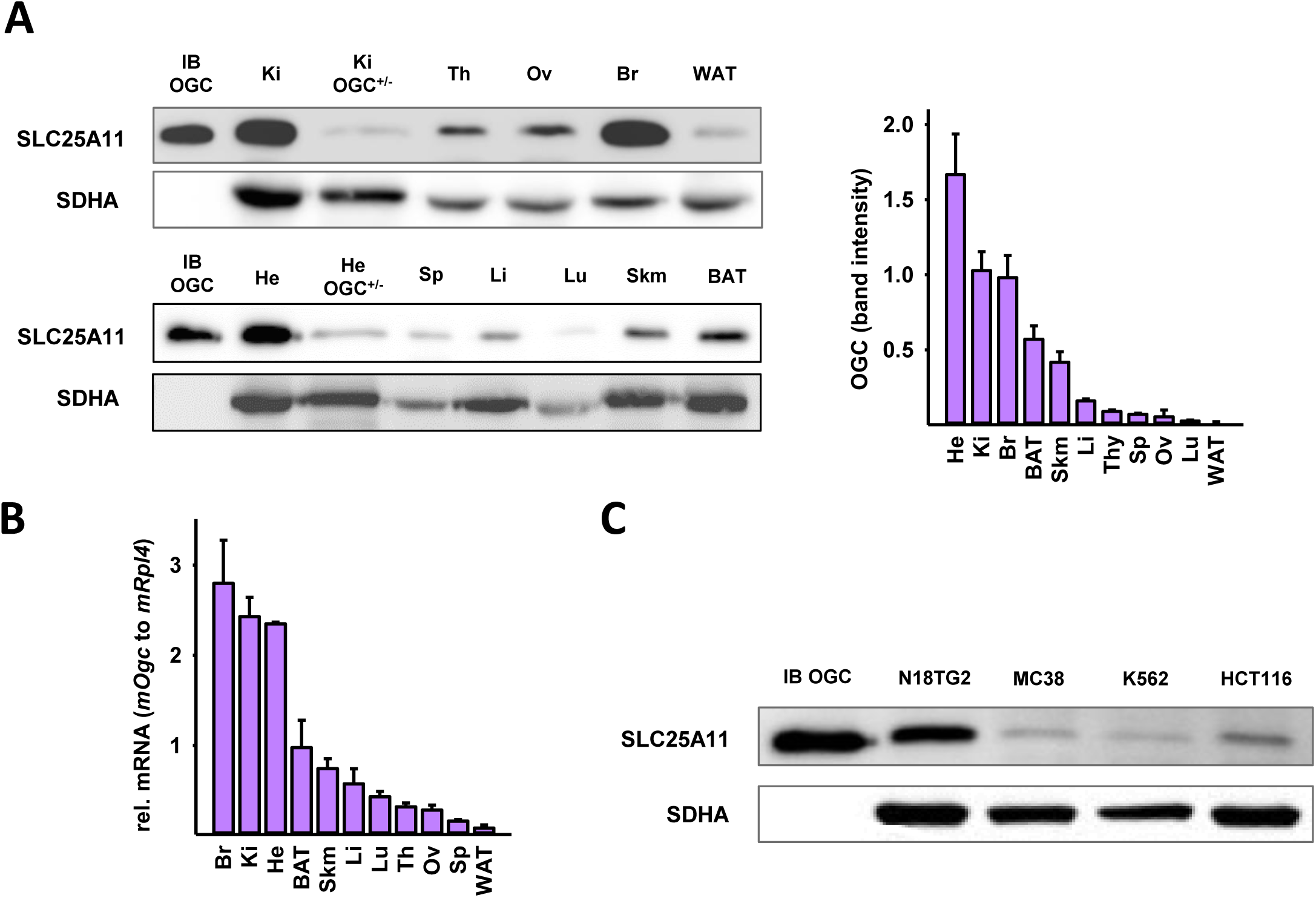
Detection of OGC (SLC25A11) in mice and cancer cell lines. (A) Representative Western blot of OGC (SLC25A11) levels in mouse tissues. Tissues are labeled as follows: kidney (Ki), thymus (Th), ovary (Ov), brain (Br), white adipose tissue (WAT), heart (He), spleen (Sp), liver (Li), lung (Lu), skeletal muscle (Skm), and brown adipose tissue (BAT). The graph shows the quantification of OGC band intensity in all tissue samples, divided by 10^8^. (B) Relative mRNA levels (2^-ΔCt^) of mOGC using ribosomal protein L4 (mRPL4) as a reference gene. (C) Western blot analysis of OGC in the following cell lines: mouse neuroblastoma cell line (N18TG2), mouse colon adenocarcinoma cell line (MC38), human myelogenous leukemia cell line (K562), and human colorectal carcinoma cell line (HCT116). 40 (A) or 20 μg (C) of total protein was loaded per lane. Recombinant mouse OGC inclusion bodies (IB OGC, 2 µg) were used as positive controls, and mouse OGC heterozygous heart (He^+/-^) and kidney (Ki^+/-^) knockouts as negative controls in (A). OGC is detected at the corresponding size of 34 kDa. Succinate dehydrogenase (SDHA) was used as a mitochondrial control. Reversible Ponceau S staining loading control, full blots and anti-OGC antibody validation are shown in Fig. S1. See *Materials and Methods* for further details.

### 2. Production and reconstitution of mOGC in liposomes

OGC was the first eukaryotic protein to be expressed in *E. coli* and functionally reconstituted in proteoliposomes ^9^. The homology between OGC and other proteins of the SLC25 superfamily, such as ANT1, UCP1, and the dicarboxylate carrier is high, especially in conserved mitochondrial carrier motifs and substrate binding sites (Fig. 2). Therefore, it is possible to express and purify them by using similar protocols.

**Figure 2.**
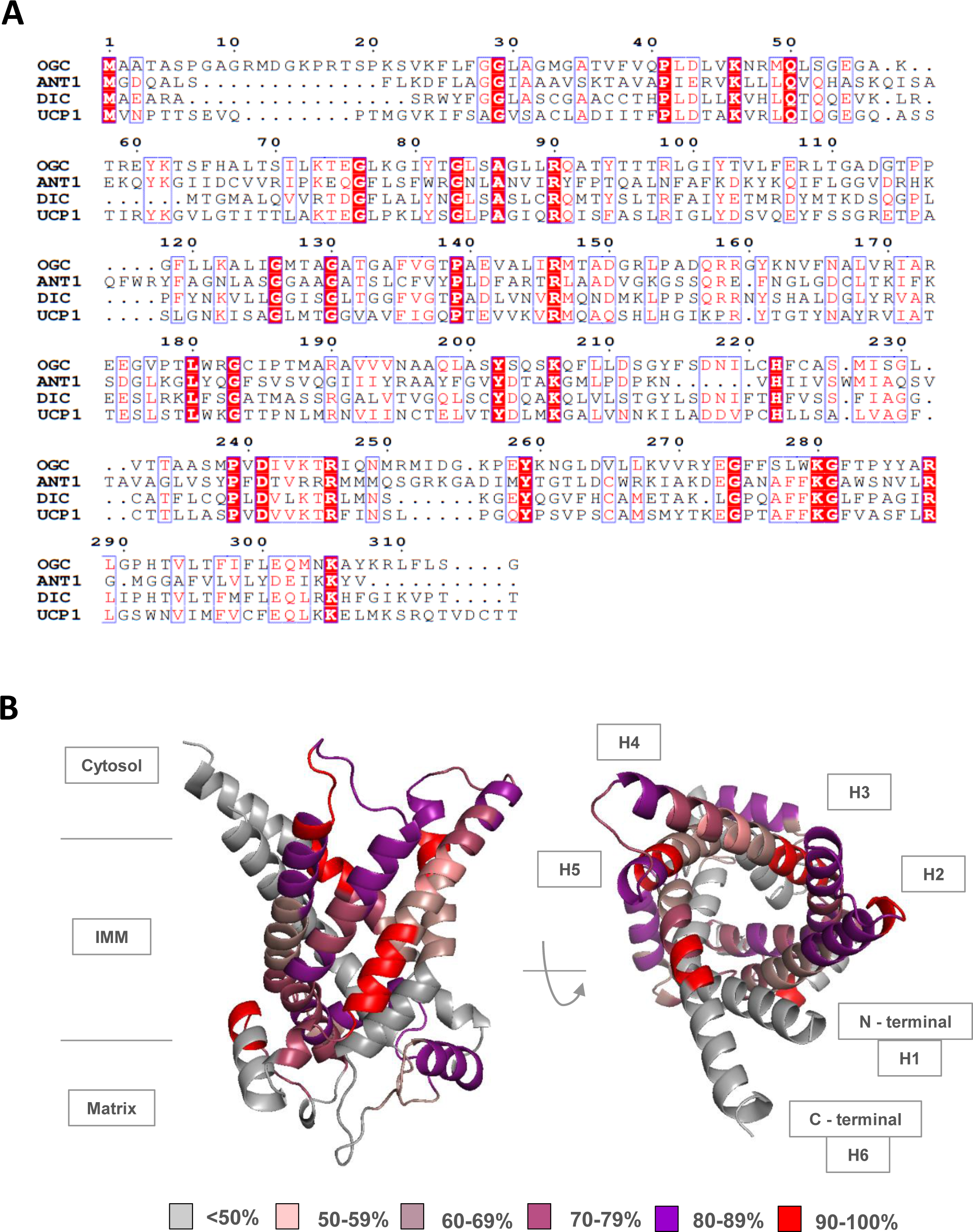
High sequence homology in substrate binding sites and signature motifs between OGC and other members of the SLC25 family. **A.** Sequence alignment of murine OGC, adenine nucleotide translocase 1 (ANT1), dicarboxylate carrier (DIC), and uncoupling protein 1 (UCP1). Generated with ClustalW and colored with ESPript. **B.** Sequence homology scores between mOGC and mANT1. Homology scores are averaged for every 5 amino acid residues based on their individual homology percentage scores calculated in the alignment. The color key is: < 50% = gray, 50-59% = green, 60-69% = lime green, 70-79% = yellow, 80-89% = orange, and 90-100% = red. The image was generated in PyMOL using the AlphaFold structure of mOGC (AF-Q9CR62-F1) as a template. Alpha helices are numbered H1-H6. The first 20 amino acids of the N-terminus are not shown here for simplicity.

In this study, we adapted previously established protocols to produce murine OGC (mOGC) in *E. coli* inclusion bodies ^9,17,22^. After purification and reconstitution into proteoliposomes (see *Materials and Methods*), it was present as a dimer (Fig. 3A). Early cross-linking studies suggested possible cysteine-linked dimerization of OGC in detergent ^23^, although the issue remained controversial ^24^. The observed dimerization may be a result of aggregation under SDS-PAGE conditions ^25^.

**Figure 3.**
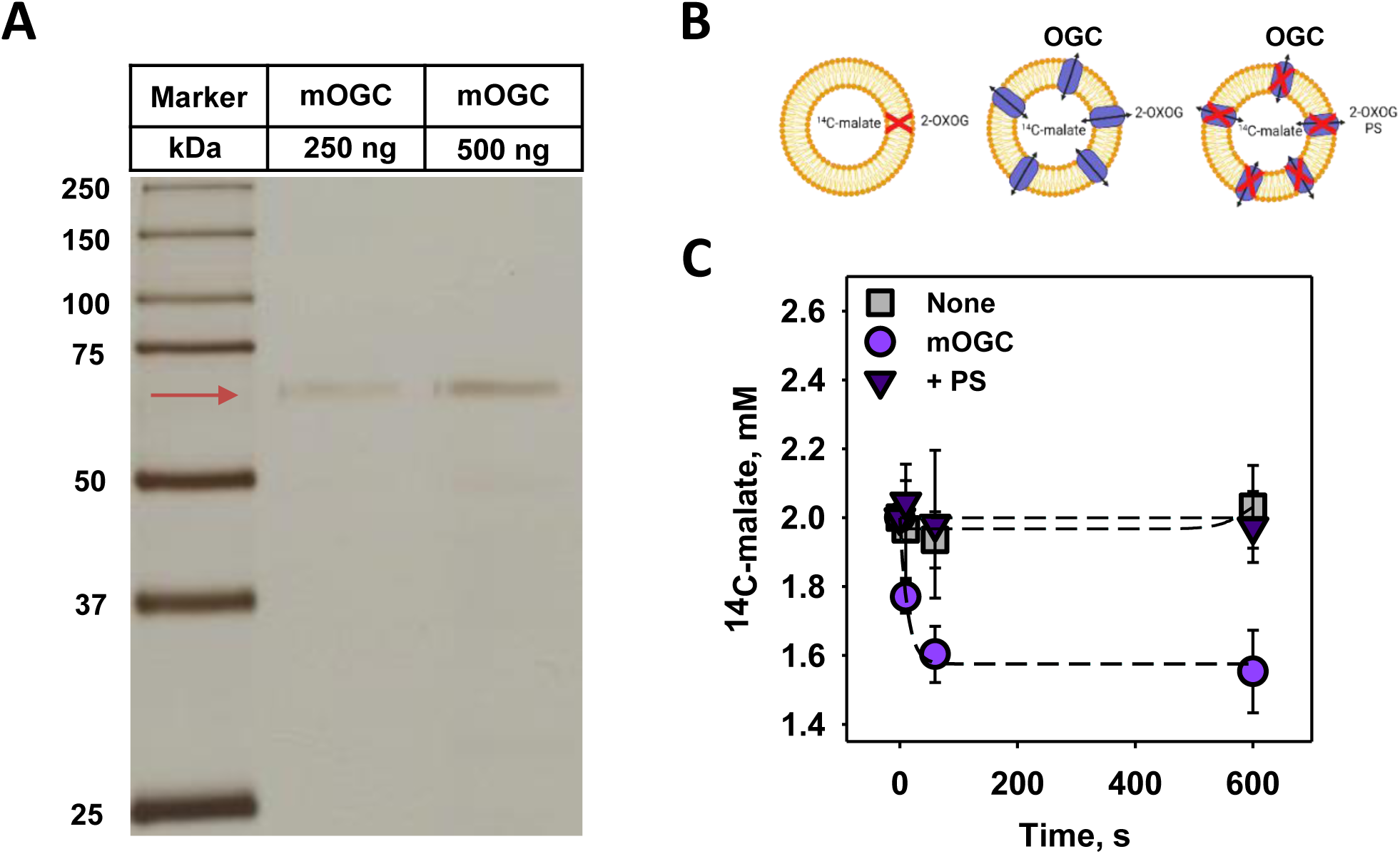
Reconstituted mOGC transports ^14^C-malate for 2-oxoglutarate. **A.** Representative silver staining of mOGC reconstituted into proteoliposomes. 250 and 500 ng of proteoliposomes were loaded in separate lanes on a 15% acrylamide gel, separated by SDS-PAGE and visualized by silver staining. Reconstituted mOGC appears at the expected dimer size (∼69 kDa, red arrow). Precision Plus Protein Dual Color Standard was loaded as a molecular weight marker. **B.** The scheme shows the three experimental setups: empty liposomes (left), active transport with mOGC (middle), and inhibition of transport by phenylsuccinate (PS, right). **C.** Decrease in ^14^C-malate concentration upon initiation of 2-oxoglutarate/malate exchange in proteoliposomes containing reconstituted mOGC (circles). The transport was not measured in empty liposomes (squares) and was completely inhibited by the addition of 20 mM of the substrate analog PS (triangles). For all measurements, membranes were prepared DOPC:DOPE:CL (45:45:10 mol %). Lipid and protein concentrations were 1.5 mg/ml and 4 µg/(mg of lipid), respectively. The buffer solution consisted of 50 mM Na_2_SO_4_, 10 mM Tris, 10 mM MES and 0.6 mM EGTA at pH = 7.34 and T = 32°C. 100 nm proteoliposomes were filled with 2 mM ^14^C-malate and transport was initiated by the addition of 2 mM 2-oxoglutarate from the outside. ^14^C-malate, oxoglutarate and PS were dissolved in buffer (pH = 7.34). Data are mean ± SD of at least three independent experiments.

### 3. OGC transports ^14^C-malate for 2-oxoglutarate

To confirm the functionality of OGC and its correct refolding in proteoliposomes, we performed substrate transport exchange measurements using ^14^C-malate and 2-oxoglutarate (Fig. 3B). Fig. 3C shows the decrease in ^14^C-malate concentration in proteoliposomes reconstituted with mOGC upon the addition of 2-oxoglutarate. The transport rate, τ, was calculated from the decrease in ^14^C-malate concentration over time, normalized to the protein concentration. The determined τ = 47.24 µmol/min/mg is in a good agreement with the previously published results for OGC, isolated or produced in different expression systems (Supplementary Table 1). Transport was completely inhibited by phenylsuccinate (PS), a substrate analog and known inhibitor of the transport function of OGC ^26^.

### 4. Fatty acid and 2,4-dinitrophenol-mediated proton transport is facilitated by OGC in planar lipid bilayers

Next, we tested whether OGC contributes to the proton conductance in pure lipid membranes composed of DOPC, DOPE and cardiolipin (45:45:10 mol%) in the model system ^27^. The specific membrane conductance (G_m_) of protein-free planar lipid bilayers (G_m_ = 8.75 ± 3.2 nS/cm^2^) is comparable to the conductance of membranes reconstituted with OGC alone (G_m_ = 9.23 ± 3 nS/cm^2^). The G_m_ of bilayers containing arachidonic acid (AA, G_m_ = 34.1 ± 7.1 nS/cm^2^) or DNP (G_m_ = 46.8 ± 6.1 nS/cm^2^) was significantly increased in the presence of OGC (G_m_ = 92.5 ± 16.5 nS/cm^2^ for AA, G_m_ = 69.6 ± 7.5 nS/cm^2^ for DNP) (Fig. 4A, ,fig. S2).

**Figure 4.**
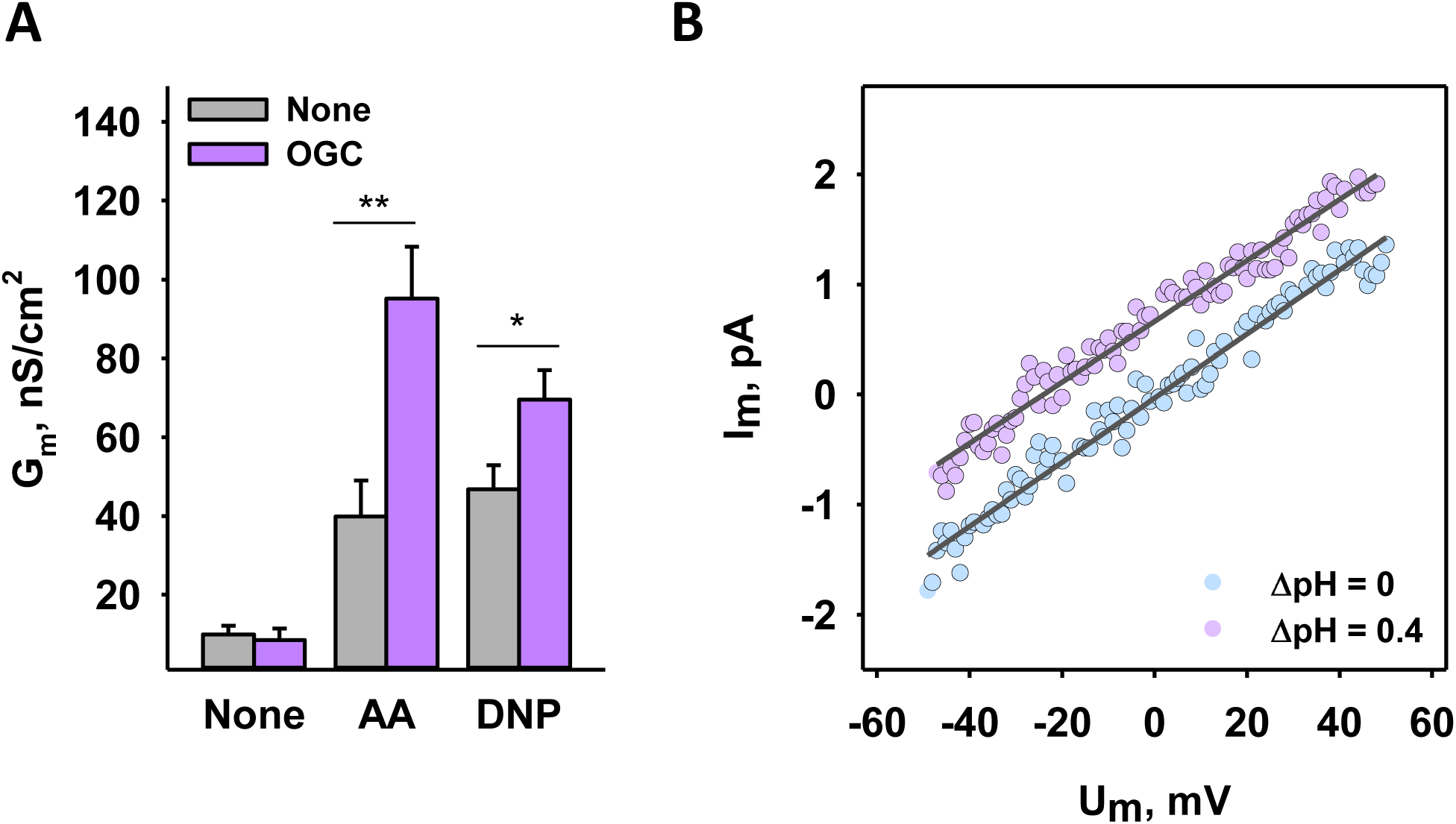
OGC-mediated proton conductance in the presence of AA or DNP in planar lipid bilayers.z. **A.** Increase in total membrane conductance (G_m_) in the presence of 15 mol% arachidonic acid (AA) or 50 µM DNP without (gray) and with (purple) OGC. **B.** Current-voltage (I/V) recordings of lipid bilayer membranes reconstituted with OGC in the presence and absence of a transmembrane pH gradient of 0.4 pH units. For all measurements, membranes were made of DOPC:DOPE:CL (45:45:10 mol%). Lipid and protein concentrations were 1.5 mg/ml and 4 µg/(mg of lipid), respectively. The buffer solution consisted of 50 mM Na_2_SO_4_, 10 mM Tris, 10 mM MES and 0.6 mM EGTA at pH = 7.34 and T = 32°C. DNP and ATP were dissolved in DMSO and buffer (pH = 7.34), respectively. Data are mean ± SD of at least three independent experiments.

To confirm that OGC facilitates proton transport in the presence of AA, and not other ions present in the buffer, we measured current-voltage characteristics in the presence and absence of a transmembrane pH gradient of 0.4 (ΔpH = 0.4) (Fig. 4B). Solutions on both the *cis* and *trans* sides of the planar lipid bilayer had the same concentrations of all ions except H^+^ and OH^-^, similar ionic strength, and similar osmolarity. The pH values on the *cis* and *trans* sides were 7.34 and 7.74, the latter being adjusted with 2.39 mM of Tris dissolved in water (pH = 7.34) after bilayer membrane formation. In this case, the experimentally obtained shift of the reversal potential, ΔV_0_, is equal to the theoretical H^+^ Nernst potential (Ψ_N_) at ΔpH = 0.4. The shift of intersection points of I/V curves with the x-axis, for ΔpH = 0 and ΔpH = 0.4, resulted in V_0_ = 25.8 ± 6.8 mV (Fig. 4B). We then calculated the transfer number of H^+^ and OH^-^ ions across the membrane (Equation 1),

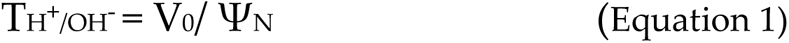

where Ψ_N_ is the theoretical value of H^+^ Nernst potential at ΔpH = 0.4 (23.9 mV). T_H+/OH-_ was 1.08 ± 0.3, which confirms that the observed increase in G_m_ is exclusively due to the transport of protons.

### 5. OGC mediates proton transport in the presence of FA with different structures

To understand whether OGC-mediated proton transport correlates with the FA saturation degree we measured the G_m_ of OGC-reconstituted lipid bilayers in the presence of FAs with increasing number of double bonds - arachidic acid (20-0), cis-11-eicosaenoic acid (20-1), cis-11,14-eicosadienoic acid (20-2), cis-8,11,14-trienoic acid (20-3), and AA (20-4) (Fig. 5). Our results show that the G_m_ is inversely correlated with the saturation degree of FA, similar to what has been observed in UCP1, UCP2, and _ANT1 28,29._

**Figure 5.**
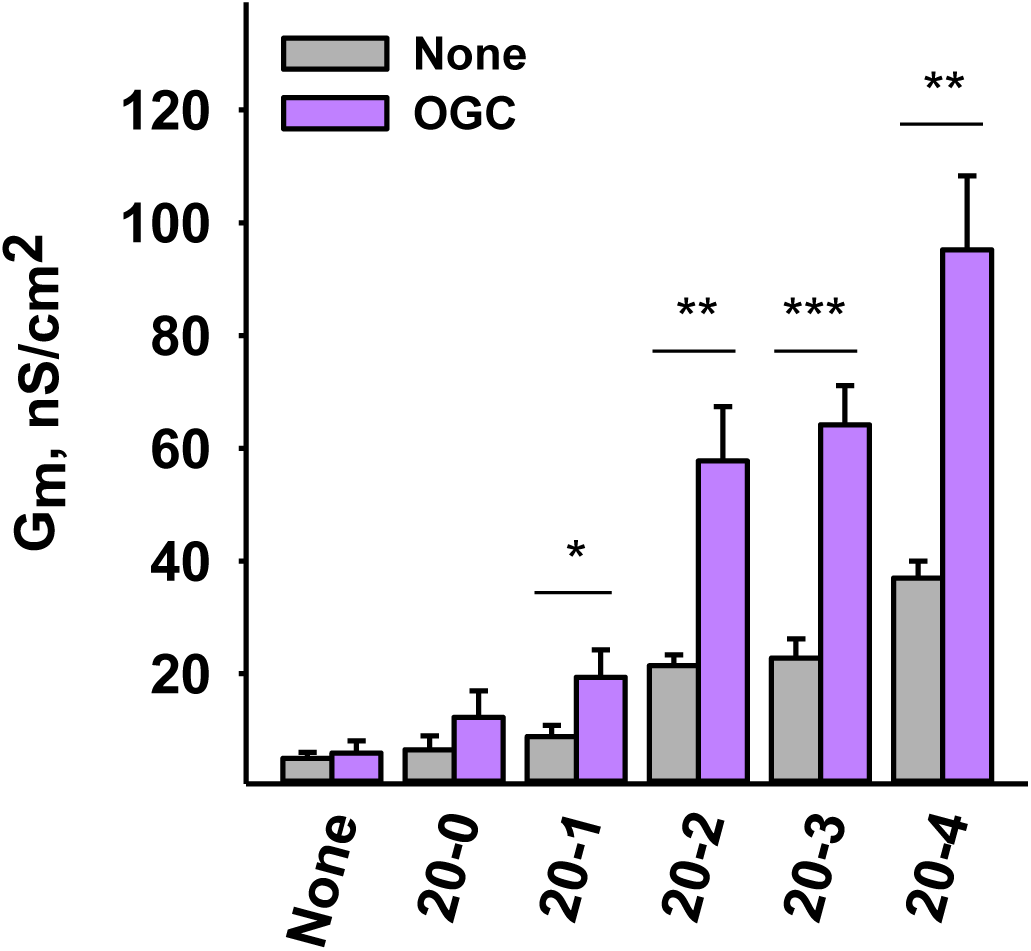
The increase in OGC-mediated proton transport directly correlates with the increase in the number of unsaturated bonds of FAs. Increase in G_m_ in the presence of arachidic acid (20-0), cis-11-eicosaenoic acid (20-1), cis-11,14-eicosadienoic acid (20-2), cis- 8,11,14-trienoic acid (20-3), and AA (20-4). The first number represents the number of carbon atoms in the FA chain; the second number represents the number of unsaturated bonds. The experimental conditions were similar to hose in Figure 4.

### 6. The activation of OGC by AA can be inhibited by ATP, its substrates, and substrate analog

We further tested whether ATP could inhibit OGC-enhanced proton transport in the presence of AA. Fig. 6A and Fig. S3A show that ATP can inhibit OGC-mediated proton conductance by 75%, indicating a possible common binding site for ATP and AA. Next, we tested whether the main substrates transported by OGC, 2-oxoglutarate and malate, show a similar effect. Fig. 6B and Fig. S3B show that the rate of AA-mediated proton transport inhibition in the presence of OGC depends on the substrate concentration with IC50s of (0.06 ± 0.01) mM and (0.1 ± 0.06) mM for 2-oxoglutarate and malate, respectively. This is consistent with the substrate transport function of OGC, and its highest affinity for 2-oxoglutarate ^9^. We then found that the IC50 of PS was (1 ± 0.11) mM (Fig. 6C, Fig. S3C), higher than both substrates tested. None of the tested compounds altered the G_m_ values when only OGC or AA were present in the system (Fig. S3D), confirming that there is no unspecific proton leak. Taken together, these results may indicate a common binding site, or a shared part of the binding site in OGC for AA, ATP, 2-oxoglutarate, malate, and PS.

**Figure 6.**
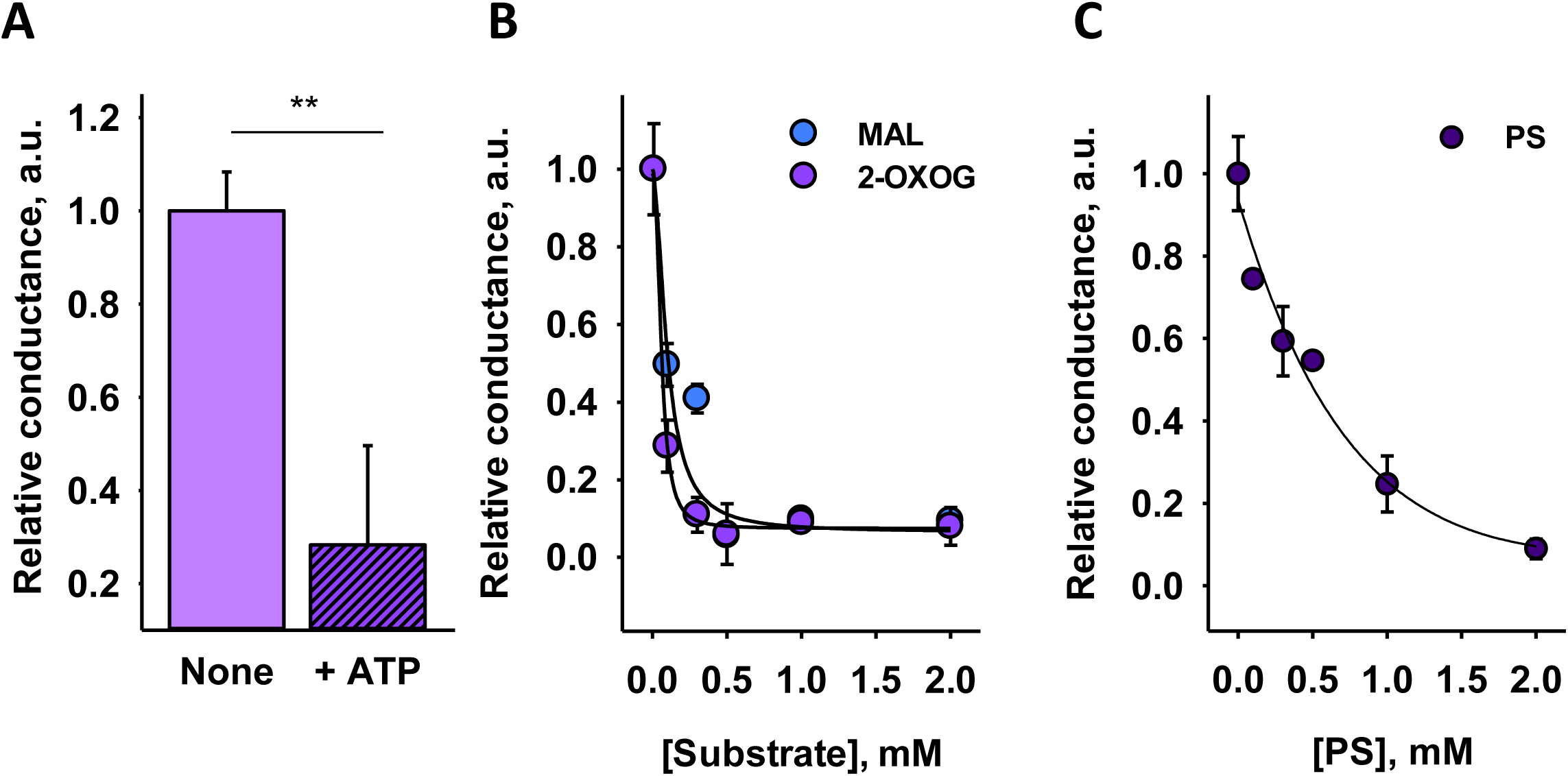
Inhibition of OGC-mediated proton transport in the presence of AA by ATP, 2-oxoglutarate (2-OXOG), malate (MAL) and phenylsuccinate (PS). Relative conductance of OGC activated by 15 mol% AA and inhibited by 4 mM ATP (**A**), 2-OXOG or MAL (**B**), or PS (**C**). Relative conductance describes the ratio between the total membrane conductance in the presence and absence of the protein with respect to the membrane conductance measured in the presence of lipid and AA only (see *Materials and Methods*). The curves in B) and C) were fitted (least squares method) with a 4-parameter sigmoidal function and IC50 values of (0.06 ± 0.01) mM for 2-OXOG, (0.1 ± 0.06) mM for MAL, and (1 ± 0.11) mM for PS were obtained. ATP, 2-OXOG and MAL were dissolved in buffer (pH = 7.34). PS was dissolved in DMSO. Other experimental conditions were similar to those described in Figure 4.

### 7. R90 is involved in the binding of FAs, ATP, and substrates

Residues R90, Y94, R98, R190, and R288 were identified as critical for substrate transport of OGC by substitution with cysteine and leucine ^30–32^. Using site-directed mutagenesis, we produced mOGC-R90S (Fig. 7A) and compared the specific membrane conductance of the bilayer membranes reconstituted with the wt and mutant proteins. Figure 7B shows that G_m_ is 50% lower in the presence of mOGC-R90S (63.28 ± 5.12 nS/cm^2^ versus 92.47 ± 16.49 nS/cm^2^ measured for mOGC, relative to the AA-induced G_m_). Furthermore, mutation of R90 to serine completely abolished the ability of ATP to inhibit mOGC. We also tested how efficiently mOGC-R90S can be inhibited by malate and 2-oxoglutarate. Figures 7C and S4 show that 1 mM of 2-oxoglutarate or malate inhibited the activated OGC by 0% and 20%, respectively, whereas 2 mM of 2-oxoglutarate or malate inhibited the protein by 19% and 74%, respectively. These results suggest that R90 may be more important in the binding of 2-oxoglutarate than malate in the proteińs cavity, or that malate competes more efficiently with AA for the rest of the binding site.

**Figure 7.**
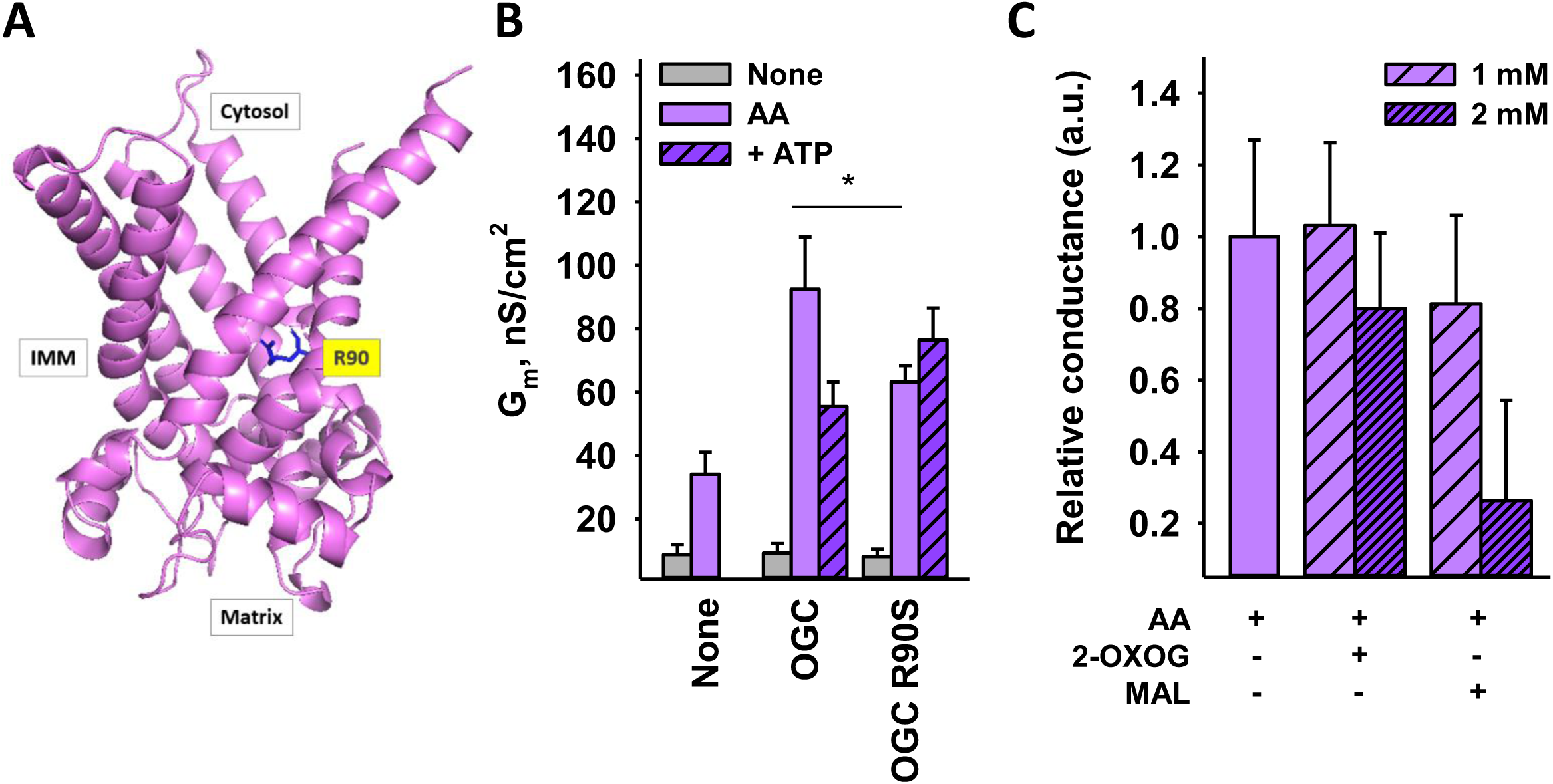
Substrate-binding residue R90 is involved in uncoupling. **A.** Side view of OGC with R90 shown in blue licorice and labeled in yellow. The image was generated in PyMOL using the AlphaFold structure of mOGC (AF-Q9CR62-F1) as a template. The N-terminal sequence has been truncated for simplicity. **B.** Comparison of total membrane conductance (G_m_) between OGC and OGC R90S in the presence of 15 mol% AA (purple) or 4 mM ATP (purple pattern). G_m_ increment mediated by OGC R90S was inhibited by 4 mM of ATP (purple pattern). **C.** Relative G_m_ measured in the presence of different 2-OXOG or MAL concentrations. Other experimental conditions were similar to those described in Figure 4.

## Discussion

Our results show that OGC can participate in uncoupling under similar conditions as ANT1 and UCP1-3, and that proton transport is enhanced only in the presence of the artificial uncoupler DNP or long-chain FAs ^17^. Proton transport facilitated by OGC increased with the degree of unsaturation of the FA, as has been shown for other carriers ^28,29^. Furthermore, ATP can competitively inhibit the activity of ANT1, UCP1-UCP3 and OGC in the presence of arachidonic acid, suggesting a common binding site for nucleotides. We identified R90 as one of the residues involved in the binding of AA and ATP to OGC (Fig. 7). This amino acid is a part of the conserved common substrate binding site of the SLC25 family and is a homolog of R79 in ANT1 and R84 in UCP1, which are critical for the binding of nucleotides, FAs, and DNP ^18,33–37^. It also has a homolog in UCP2 (R88) and dicarboxylate carrier (R69). Since these proteins are involved in both substrate transport and FA-mediated proton transport, we propose that OGC is a dual-function protein whose substrate binding site is part of the interaction site with negatively charged uncouplers such as FAs or DNPs (Fig. 8).

**Figure 8.**
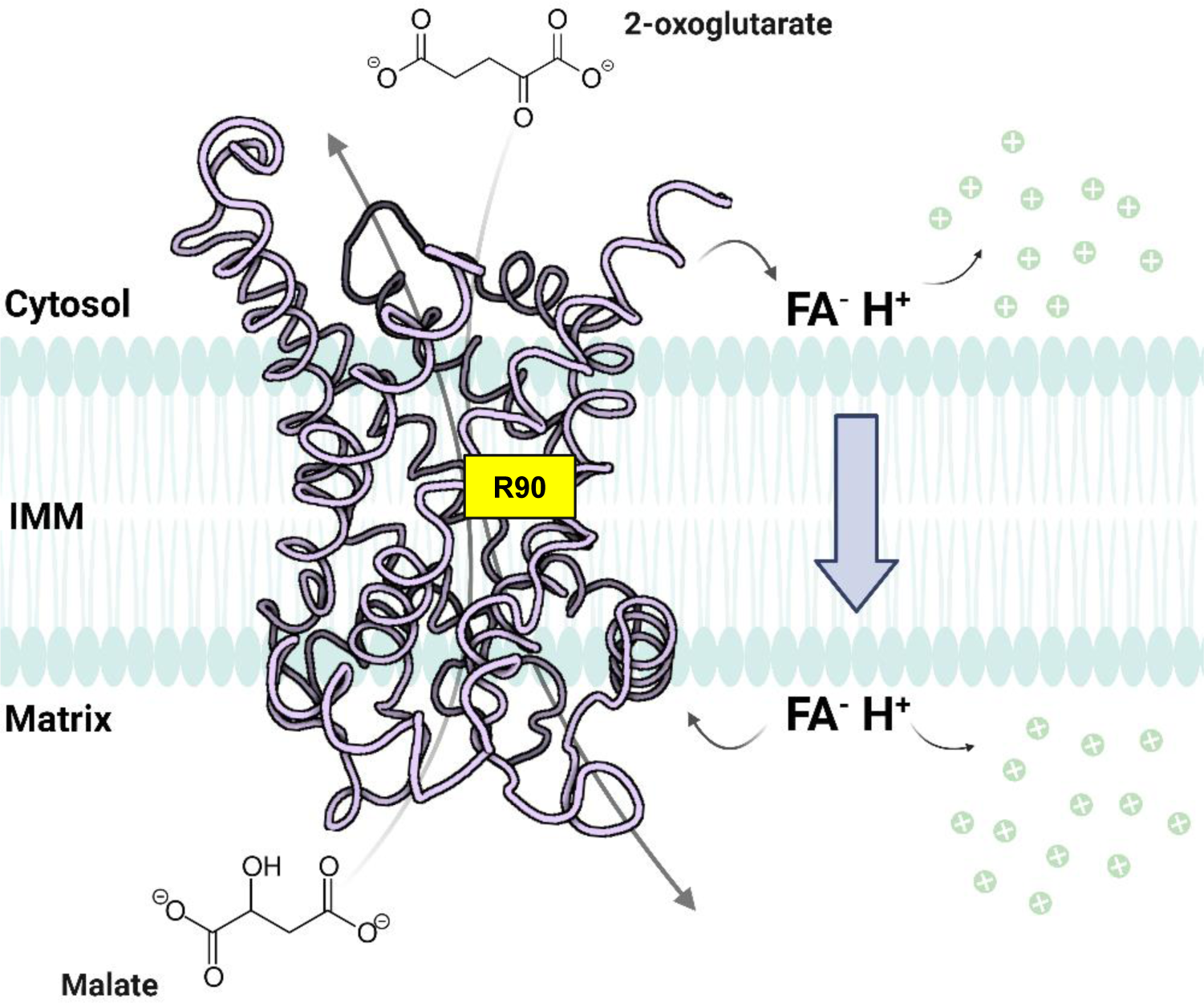
OGC catalyzes substrate transport and participates in FA-mediated uncoupling. R90 is a critical part of both the substrate and FA binding sites. Image is generated in BioRender using the AlphaFold structure of mOGC (AF-Q9CR62-F1) as a template. The N-terminal sequence has been truncated for simplicity.

Unlike the aspartate/glutamate carrier, which has a confirmed dimerization interface via the regulatory domain, all other known SLC25 family proteins apparently exist and function as monomers ^24^. Previous results showing otherwise ^22,38–41^ have been disputed for their use of harsh detergents or lack of proper understanding of the experimental data ^42^. However, the oligomeric state of OGC remained uncertain. Cross-linking studies strongly suggested dimerization in the presence of detergent, with the dimer formed with a single disulfide bond between two C184 residues across the subunits. This bond was present regardless of protein concentration, strongly contradicting the hypothesis of the random aggregation ^23,40^. Despite these findings, the dimerization of OGC in the membrane has not been further demonstrated and therefore remains questionable ^24^. While assessing the surface residues of OGC, it was found that none of them could constitute a conserved homo-dimerization interface. Some residues were identified as potential candidates for specific recognition, but these are not conserved among SLC25 members. It is considered unlikely that distinct homo-dimerization interfaces exist for each carrier ^43^. However, this does not exclude the possibility of dimerization, especially since the formation of a disulfide bond by cysteine 184 was not considered in this case.

A possible explanation for thermogenesis-independent uncoupling would be mild uncoupling or lowering of MMP to prevent the generation of damaging ROS ^44,45^ . During oxidative stress, the activity of phospholipid hydrolysis catalyzed by phospholipase A significantly increases ^46^. As a result, the bloodstream concentration of FAs increases in pathological conditions such as obesity and type II diabetes ^47^. Besides free FAs, oxidative stress leads to the formation of lipid hydroperoxides and reactive aldehydes, which can modify and activate UCPs ^48,49^. Under these conditions, OGC is likely involved in FA-mediated proton transport.

However, we found that 2-oxoglutarate is a competitive inhibitor of OGC-mediated proton leak, with an EC50 value of 60 µM (Fig. 6B). Considering that human plasma, liver and brain contain 10-60 µM ^50^, 150-300 µM ^51^ and 600-800 µM ^52^ of 2-oxoglutarate, respectively, it seems that the cavity of OGC is permanently occupied and not available to bind FAs. Yet, it has been observed that 2-oxoglutarate levels are significantly reduced when mitochondrial function is impaired or damaged by oxidative stress, such as in diabetes ^53^ and breast cancer ^54^. Under conditions that promote elevated levels of FAs and reactive aldehydes, it is plausible that OGC participates in proton transport to prevent further damage. It remains to be unravelled whether OGC alone contributes to significant proton leak *in vivo*. Unlike UCP1 in BAT under cold acclimation conditions, individual proteins such as OGC, ANT1, UCP2, or UCP3 may not induce a substantial amount of proton transport. However, their collective uncoupling function could lead to a beneficial and transient decrease in MMP, mitigating further oxidative damage without the need for upregulation of protein expression.

In summary, by reconstituting OGC as the only protein in the lipid bilayer membrane, we have proven its role in facilitating proton transport in the presence of FAs and DNP. ATP, OGC substrates, and PS inhibited this effect, indicating a competition for the same binding site. We identified residue R90 which is involved in both uncoupling and substrate transport in OGC and may play a critical role in the conserved mechanism of uncoupling throughout the SLC25 family.

## Materials and Methods

### 1. Chemicals

1,2-dioleoyl-sn-glycero-3-phosphotidylcholine (DOPC, #SLCF9767), 1,2-dioleoyl-sn-glycero-3-phosphoethanolamine (DOPE, #SLCB7462), cardiolipin (CL, #SLCC2621), adenosine 5′-triphosphate (ATP, #SLBZ3783), α-ketoglutaric acid (2-oxoglutaric acid, #BCCC4294), phenylsuccinic acid (#MKBS0493V), L-malic acid (#SLCD6882), N-laurylsarcosine (#L5125), Triton X-114 (#MKCL3391), Triton X-100 (#STBJ5677), dithiothreitol (DTT, #BCCG5712), 2,4-dinitrophenol (D198501), dimethylsulfoxide (DMSO, #276855), bovine serum albumin (BSA, #SLCM9403), arachidic acid (Ara, #0000145297), cis-11-eicosaenoic acid (20-1, #MKCP4439), cis-11,14-eicosadienoic acid (20-2, #SLCL2548), cis-8,11,14-trienoic acid (#SLCG0821), protease inhibitor cocktail (#P8340), bromophenol blue (#BCBF8233V), hexane (#296090), hexadecane (#296317), and isopropanol (#I9516) were purchased from Sigma-Aldrich (Vienna, Austria). Sodium sulfate (Na_2_SO_4_, #8560.3), sodium chloride (NaCl, #9265.1), potassium chloride (KCl, #6781.3), 2-(N-morpholino) ethanesulfonic acid (MES, #4256.2), tris(hydroxymethyl)-aminomethane (Tris, #4855.2), ethylenediaminetetraacetic acid (EDTA, #8043.2), ethylene glycol-bis(β-aminoethyl ether)-N,N,N′,N′-tetraacetic acid (EGTA, #8043.1), isopropyl β-D-1-thiogalactopyranoside (IPTG, #2316.3), chloramphenicol (#3886.2), kanamycin (#T832.1), β-mercaptoethanol (#4227.3), glycerine (#3783.1), ethanol (#T913.1), desoxycholic acid sodium salt (#3484.2), sodium dodecyl sulfate (SDS, #0183.3), agarose (#3810.2), and Ponceau S staining solution (#5938.2) were purchased from Carl Roth GmbH & Co. K.G. (Karlsruhe Germany). Chloroform (#AE 54.1) was obtained from either Carl Roth GmbH & Co. K.G. (Karlsruhe Germany) or PanReac AppliChem (UN1888, Darmstadt, Germany). We purchased arachidonic acid (AA, #10-2004-7) from Larodan (Solna, Sweden), n-octylpolyoxyethylene (#1000013726) from BACHEM (Bubendorf, Switzerland), and hydroxyapatite (#130-0420), Bio-Beads SM-2 (#152-3920) and the enhanced chemiluminescence (ECL) western blotting reagent (#170-50001) from Bio-Rad Laboratories (Hercules, CA, USA). Dulbecco’s phosphate-buffered saline (DPBS, #14190144) was obtained from Thermo Fisher Scientific, Waltham, MA, USA), PhosSTOP™ (#59124600) from Roche Diagnostics (Mannheim, Germany), and nuclease-free water (#7732-18-5) from VWR (Vienna, Austria). ^14^C-malic acid was purchased either from Perkin Elmer (Waltham, MA, USA) (#2625350) or from Hartmann Analytic (Braunschweig, Germany) (ARC 0771). The Ultima-Gold^TM^ scintillation liquid (#7722011) was purchased from Perkin Elmer (Waltham, MA, USA) and Sephadex^TM^ G-50 (#10297028) from Cytiva Sweden AB (Uppsala, Sweden).

### 2. Animals and protein sample preparation

Two-month-old female C57BL/6 wild-type mice used in this study were kept under standardized laboratory conditions (12:12 h light/dark cycle, room temperature (24°C), food and water *ad libitum*) and sacrificed by CO_2_ asphyxiation. Mouse organ and tissue samples were pooled from at least 5 mice to obtain enough protein. They were homogenized with a mixer mill (MM200, Retsch, Germany) in RIPA buffer (50 mM Tris, 150 mM NaCl, 1% desoxycholic acid sodium salt, 1 mM EDTA, 1% Triton X-100, 0.1% SDS) supplemented with a protease inhibitor cocktail. After 30 min of incubation on ice, the lysates were centrifuged 2 × 10 min at 2500 x g. The supernatants were collected, aliquoted, and stored frozen at -20°C.

Total protein isolation from cancer cells was performed as described in ^55^. In brief, the cells were washed twice in ice-cold DPBS and centrifuged at 300 x g for 5 min. Pellets were snap frozen and sonicated in RIPA buffer supplemented with a protease inhibitor cocktail and PhosSTOP™. Lysates were centrifuged 2 x 10 min at 2500 x g, and the collected supernatants were aliquoted and stored at -20°C. Total protein concentrations of tissue and cancer cell samples were determined using the Pierce BCA Protein Assay Kit (#RG235622, Thermo Fisher Scientific, Waltham, MA, USA). Further steps were performed as described for protein isolation from mouse organ tissues.

### 3. Western blot analysis

Western blotting was adapted to the previously published protocol ^56^. In brief, total protein was separated on 15% SDS-PAGE gels and transferred to nitrocellulose membranes. Reversible Ponceau S staining was used as a loading control (Fig. S1A). After blocking the membranes in 2% BSA blocking solution at RT, they were incubated with primary antibodies against OGC (anti-SLC25A11, sc-515593, #G2016, Santa Cruz Biotechnology, Dallas, TX, USA) or succinate dehydrogenase (SDHA, ab14715, #G3365497-8, Abcam, Cambridge, UK) overnight at 4 °C. Detection was performed with the UVP ChemStudio Imaging System (Analytik Jena, Jena, Germany) using horseradish peroxidase-linked anti-mouse (#38) and anti-rabbit (#29) secondary antibodies (Cell Signaling Technology, Danvers, MA, USA), and the ECL western blotting reagent. All primary and secondary antibodies were diluted with 2% BSA block solution. The semi-quantitative analysis of western blots was done using Vision Works software version 9.1 (Analytik Jena, Jena, Germany). SDHA was used as a mitochondrial marker. The values were averaged from at least 3 different membranes and 2 biological replicates. For each biological replicate, tissues from 5 mice were pooled together to isolate sufficient protein levels.

### 4. mRNA expression analysis

RNA isolation and quantitative reverse transcription (qRT-PCR) was performed as previously described ^56^. RNA was isolated using the innuSOLV RNA Reagent (Analytik Jena, Jena, Germany) according to the guanidine isothiocyanate/phenol method. Briefly, homogenized tissue samples were incubated with 1 ml of the innuSOLV RNA reagent per 100 mg. Phase separation was done by the addition of chloroform, and RNA was precipitated with isopropanol and washed with ethanol. Air-dried RNA was dissolved in nuclease free water and quantified using the NanoDrop® UV-Vis Spectrophotometer (Thermo Fisher Scientific, Waltham, MA, USA).

Two µg of RNA were subjected to DNase digestion and reverse transcribed to cDNA using random hexamer primers and the High-Capacity cDNA Reverse Transcription Kit (both Thermo Fisher Scientific, Waltham, MA, USA). RT-qPCR was conducted on qTOWER³ 84 (Analytik Jena, Jena, Germany) at a 62°C annealing temperature using the Luna® Universal qPCR Master Mix (New England BioLabs GmbH, Frankfurt am Main, Germany) and primers for *Mus musculus* solute carrier family 25 member 11 (SLC25A11) sourced from the NCBI primer-blast [NM_024211.3]. Primers targeting both murine splice variants were used: (5’->3’ forward: GTTGTTTGAGCGCCTGACTG, reverse: CAGCTGGAAGCCGACCAT). Relative expression levels of *mOgc* were calculated by the Δ cycle threshold (Ct) to the housekeeping gene *mRpl4* (2^−[Ct(*mOgc)*−Ct(*mRpl*4)]^).

### 5. Recombinant protein production and purification

Cloning, purification, and reconstitution of murine OGC was adapted to the previously established protocols ^9,57^. In brief, pET24a expression plasmids containing selected OGC cDNA sequences were transformed into the *E. coli* Rosetta (DE3) pLysS strain (Novagen^®^, Darmstadt, Germany) and selected on kanamycin (25 µg/ml) and chloramphenicol (34 µg/mL) containing plates. Successful transformants were inoculated into the growth medium containing chloramphenicol (34 µg/mL) and grown until the optical density at 600 nm reached 0.5. Protein production was then induced with 0.5 mM IPTG, and the cells were collected by centrifugation after 3 h. Inclusion bodies containing the expressed protein were isolated via high-pressure homogenization and centrifugation. The expression of mOGC in inclusion bodies was confirmed using the Western blot analysis (Fig. S1C).

### 6. Reconstitution of OGC into liposomes

To purify and reconstitute the protein, 2 mg of inclusion bodies were solubilized in a TE/G buffer containing 2% N-lauroylsarcosine, 1.3% Triton X-114, 0.3% N-Octylpolyoxyethylene, 1 mM DTT and 2 mM phenylsuccinate at pH 7.5. The lipid mixture (DOPC:DOPE:CL; 45:45:10 mol%) was hydrated overnight and added in gradually. The mixture was concentrated and dialyzed against a buffer used in the experiments (50 mM Na_2_SO_4_, 10 mM MES, 10 mM Tris, and 0.6 mM EGTA at pH 7.34). The dialysate was centrifuged and applied to a hydroxyapatite column to remove unfolded and aggregated protein fractions. Subsequently, non-ionic detergents were removed by Bio-Beads SM-2. The final protein concentration was measured with the Micro BCA Protein Assay Kit (#OI191202, Thermo Fisher Scientific, Waltham, MA, USA). Protein purity was verified by SDS-PAGE and silver staining (Fig. 3A). Proteoliposomes were produced in independent batches. Following batch numbers were used for this study: OGC #5, #9, #11, #12, #15, #23, #24, #25.

### 7. Generation of the OGC R90S mutant

*In vitro* site-directed mutagenesis was carried out on expression plasmids containing the cDNA of mOGC as a template. The mutation was introduced with an oligonucleotide designed to alter codon Arg90 (CGC) to Ser (AGC) using a Q5^TM^ site-directed mutagenesis kit (#EO552, #EO554S, New England BioLabs GmbH, Frankfurt am Main, Germany). The successful introduction of mutations was confirmed by sequencing. Mutant OGC expression plasmids were transformed in the *E. coli* expression strain Rosetta(DE3)pLysS. The rest of the protocol was as described above for the OGC wild-type, and the batch numbers used for this study were #2, #4 and #5.

### 8. Substrate exchange rate measurements of OGC

OGC-containing proteoliposomes were filled with 2 mM ^14^C labelled-malate prior to extrusion. 1 mM of DTT was added to proteoliposomes after extrusion to prevent any free sulfhydryl group-mediated aggregation. OGC facilitated transport was initiated by adding 2 mM 2-oxoglutarate (2-oxoglutaric acid dissolved in the experimental buffer at pH 7.34) and stopped after size exclusion chromatography using Sephadex^TM^ G-50 dextran gels at corresponding times (Fig. 3B). The remaining radioactivity in proteoliposomes was measured by liquid scintillation counting (Tri-Carb 2100TR, Perkin Elmer, Waltham, MA, USA). In the case of inhibition, 20 mM phenylsuccinate (PS, phenylsuccinic acid dissolved in the experimental buffer at pH 7.34) was added to proteoliposomes prior to extrusion to account for the random orientation of OGC in the membrane. For control measurements, the same protocol was used on empty liposomes.

### 9. Formation of planar bilayer membrane and measurements of the electrical parameters

Planar lipid bilayers were formed from liposomes on the tip of the dispensable plastic pipettes as previously described ^27^. Membrane formation and bilayer quality were verified by capacitance measurements (C = 0.715 ± 0.03 µF/cm^2^). Capacitance did not depend on the presence of the protein, AA, or other substrates, which were added to the lipid phase before membrane formation. Current–voltage (I–U) characteristics were measured by a patch-clamp amplifier (EPC 10, HEKA Elektronik Dr. Schulze GmbH, Germany). Total membrane conductance (G_m_) was derived from the slope of a linear fit of the experimental data at applied voltages in the range of -50 mV to +50 mV. Lipid concentration was 1.5 mg/ml (1.875 mM) in all experiments. AA was dissolved in chloroform and added to the lipid phase before vacuuming. ATP, 2-oxoglutaric acid, L-malic acid, and phenylsuccinic acid were dissolved in the buffer solution (pH adjusted to 7.34) and DNP in DMSO. The amount of DMSO added to the measuring sample never exceeded 10 µL, which was previously shown not to alter the membrane conductance ^17^. The concentrations of each substrate used in the experiments are indicated in the figure legends. The relative membrane conductance was calculated according to Equation 2:

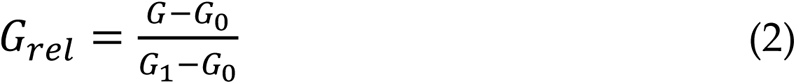

where G_0_ is the total membrane conductance of lipid membranes reconstituted with AA, G_1_ is the total membrane conductance of lipid membranes reconstituted with OGC and AA, and G is the total specific membrane conductance of lipid membranes reconstituted with OGC, AA, and/or ATP/PS/2-oxoglutarate/L-malate (Fig. 5).

### 10. Statistical analysis

Statistical analyses were performed using Sigma Plot 12.5 (Systat Software GmbH, Erkrath, Germany). Data from the electrophysiological and substrate exchange measurements are represented as mean ± standard deviation of at least three technical replicates performed on three different days. In electrophysiological measurements, each technical replicate was the mean conductance of three to ten lipid bilayer membranes formed on the same day. In substrate exchange measurements, each technical replicate stands for a new preparation of the measuring sample. Electrophysiological data were tested using the unpaired two-tailed Student’s t-test. Statistical significance was defined at p < 0.05 (*), p < 0.01 (**), p < 0.001 (***), and p < 0.0001 (****).

### Author Contributions

Conceptualization, EEP; Funding acquisition, EEP; Investigation, KZ, TT, TB, FS; Project administration, EEP; Resources, EEP; Supervision, EEP; Writing—original draft, KZ, and EEP; Writing—review & editing, KZ, TT, TB, FS, JK, EEP. All authors have read and agreed to the published version of the manuscript.

## Acknowledgments

We thank Sarah Bardakji for the excellent technical assistance, and the group of Soo-Youl Kim (National Cancer Center, Seoul, South Korea) for providing us with murine knockout samples.

## Data Availability Statement

The datasets generated and/or analyzed during this study are available from the corresponding authors upon reasonable request.

## Ethics approval statement

The Ethical Committee of the University of Veterinary Medicine, Vienna approved all animal procedures.

## Supporting information

**Figure S1.**
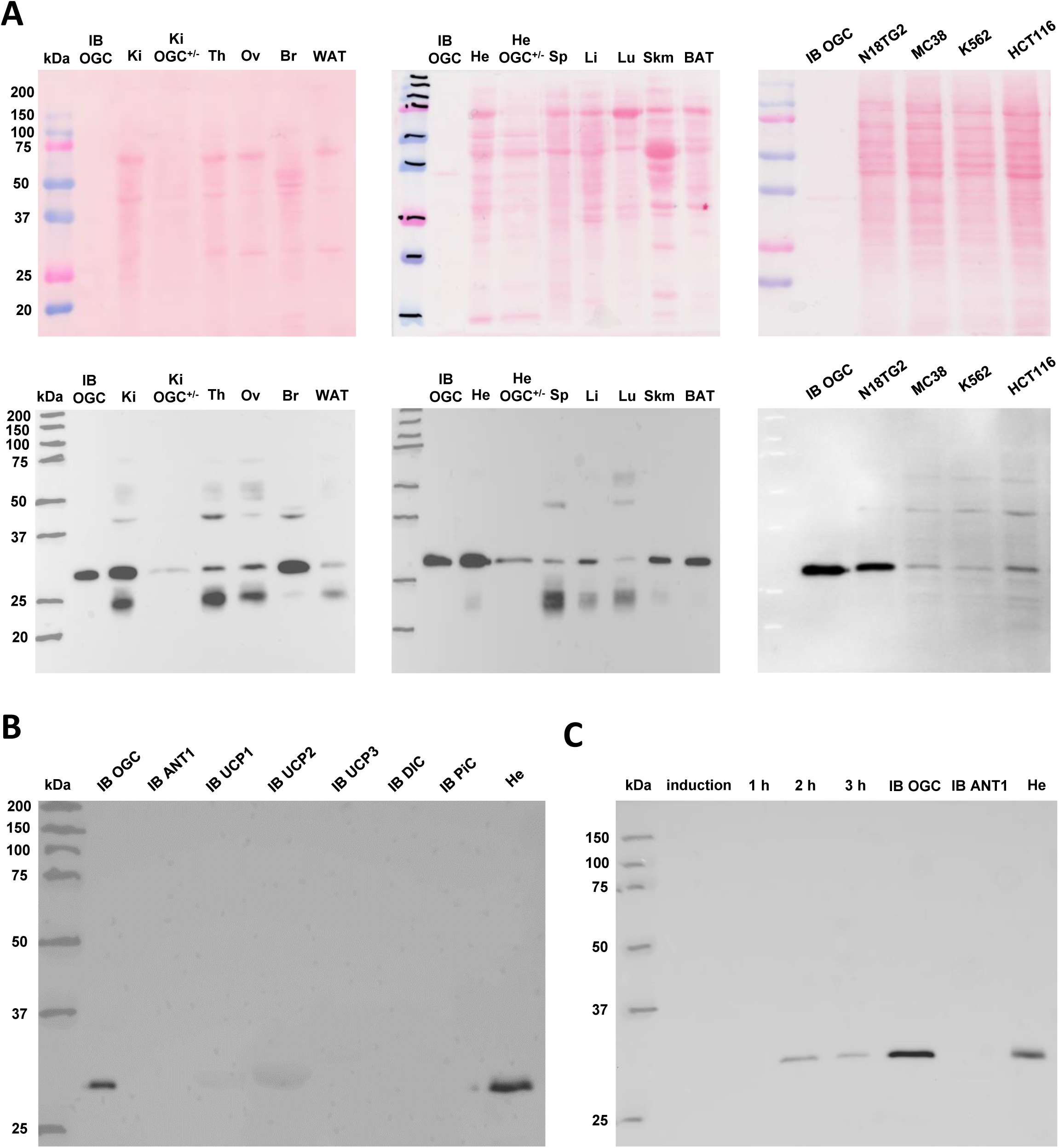
**A.** Ponceau S staining of nitrocellulose membranes (top row) and the corresponding representative blots (bottom row) developed with the anti-OGC antibody and used for the Western blot analysis in Fig. 1. The Ponceau S staining solution was applied after blotting and before the incubation with the primary anti-SLC25A11 antibody. **B.** Evaluation of the anti-SLC25A11 antibody specificity against mOGC. Inclusion bodies (IB) containing several members of the SLC25 family with a high homology to OGC were tested: adenine nucleotide translocase 1 (ANT1), uncoupling protein 1 (UCP1), uncoupling protein 2 (UCP2), uncoupling protein 3 (UCP3), dicarboxylate carrier (DIC), and phosphate carrier (PIC). 40 µg of mouse heart tissue was used as a positive control for OGC. **C.** Representative Western blot of OGC expression during protein production in *E. coli.* OGC expression was induced with 0.5 mM IPTG, and *E. coli* pellets were collected every hour. Production was stopped after 3 hours and final IB were isolated. 15 µg of total protein was loaded. Mouse heart tissue was used as a positive control and IB ANT1 as a negative control. Precision Plus Protein Dual Color Standard was loaded as a molecular weight marker.

**Figure S2.**
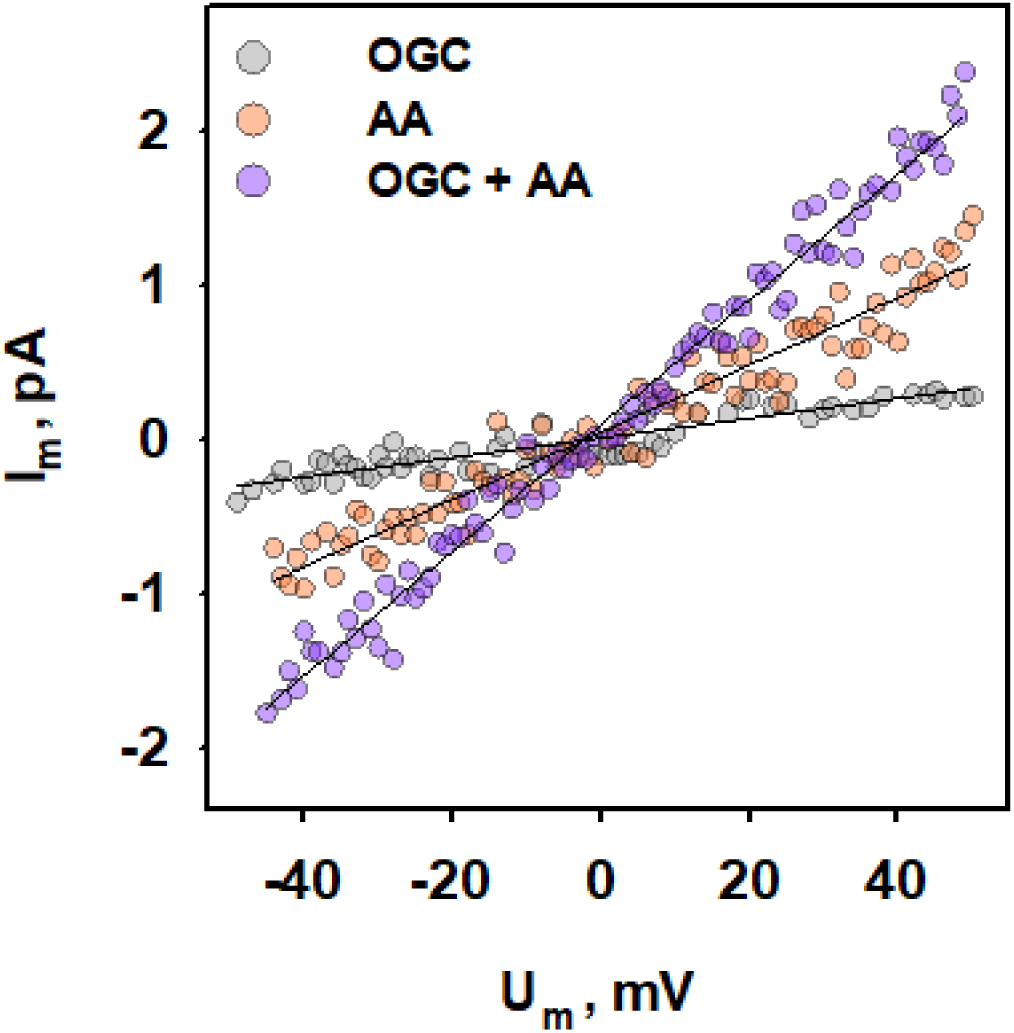
I/V characteristics of OGC-mediated AA proton transport. Representative current-voltage recordings of lipid bilayer membranes reconstituted without (grey circles) or with AA in the absence (orange squares) or presence (purple triangles) of OGC. Other experimental conditions were similar to those in Figure 4.

**Figure S3.**
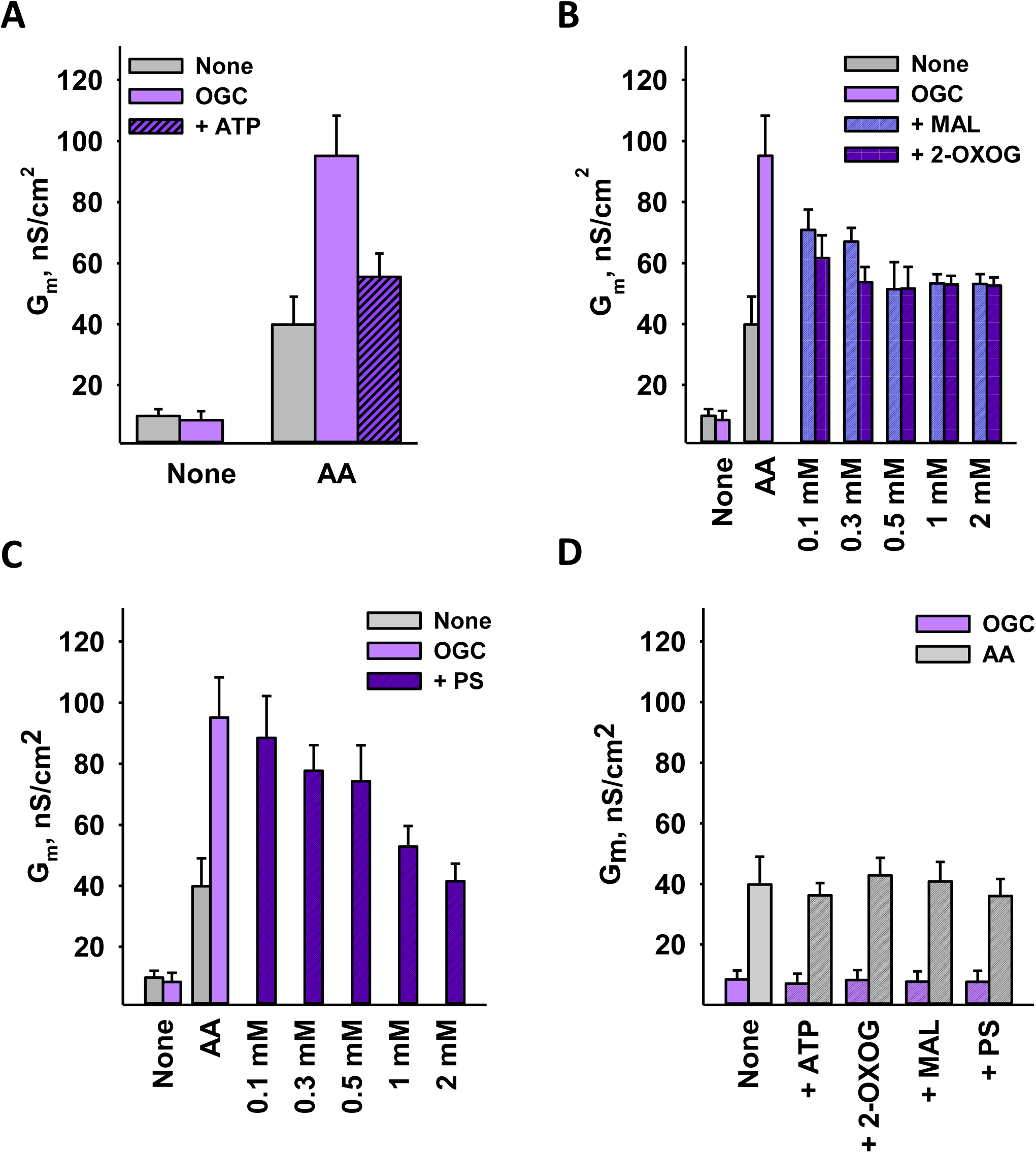
Total conductance (G_m_) of lipid bilayer membranes reconstituted with OGC and AA and inhibited with ATP (**A**), 2-oxoglutarate (2-OXOG) and malate (MAL, **B**) or phenylsuccinate (PS, **C**). These compounds had no effect on the G_m_ of lipid bilayers containing only OGC or AA (**D**). ATP, 2-OXOG and MAL were dissolved in buffer (pH = 7.34). PS was dissolved in DMSO. 2-OXOG, MAL, and PS were used at a concentration of 2 mM, and ATP was used at 4 mM, unless otherwise noted. Other experimental conditions were similar to those in Figure 4.

**Figure S4.**
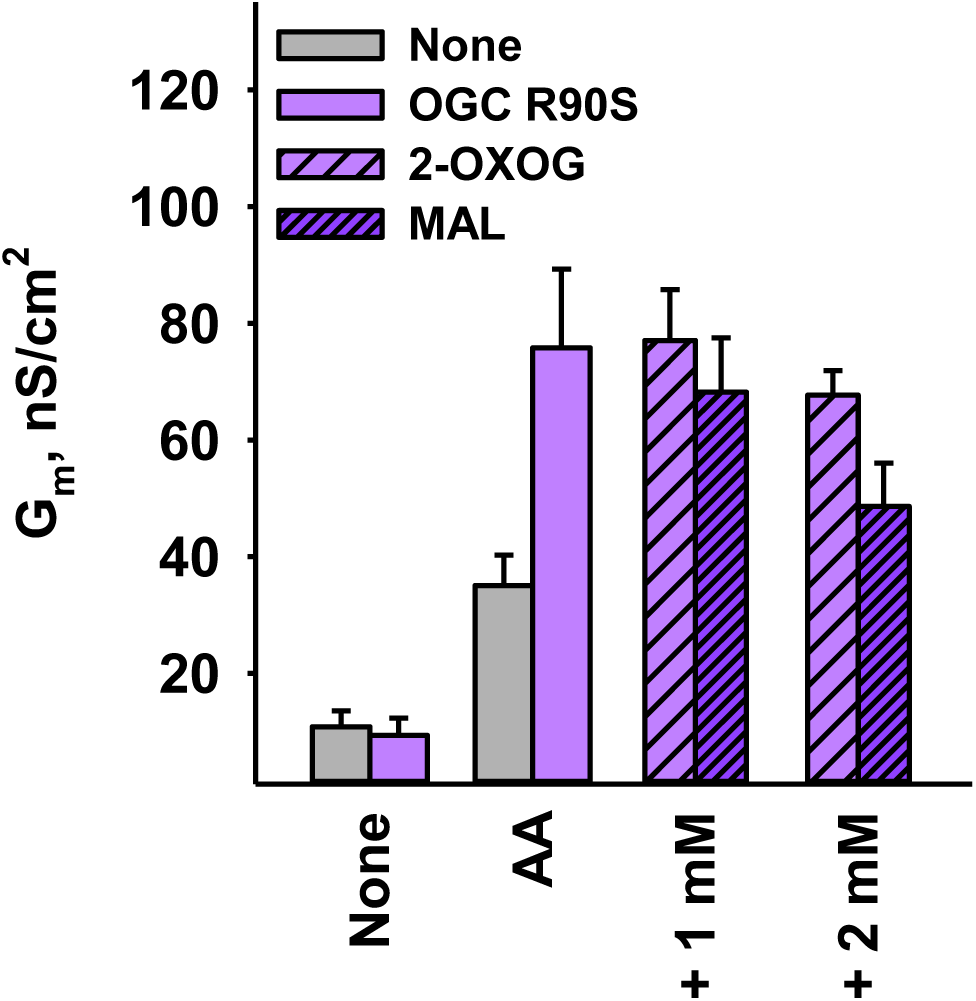
Increase in total membrane conductance (G_m_) of OGC-R90S in the presence of 15 mol% AA. The G_m_ increment was inhibited by 1 and 2 mM 2-OXOG or MAL. Other experimental conditions were similar to those in Figure 4.

**Supplementary Table 1.** Transport rates of the ^14^C-malate/2-oxoglutarate exchange.

| Protein source | $V_{\max}$ ($\mu\text{mol min}^{-1} \text{mg}^{-1}$) | Reference |
| --- | --- | --- |
| Purified from bovine heart mitochondria (2-oxoglutarate homoexchange) | 22.2 | <u>Indiveri et al., 1987</u> |
| Purified from rat liver mitochondria | 0.14 | <u>Bissaccia et al., 1988</u> |
| Purified from bovine heart mitochondria | 8-11 | <u>Indiveri et al., 1991</u> |
| Recombinant produced in *E. coli* | 0.35 | <u>Fiermonte et al., 1993</u> |
| Recombinant produced in *E. coli* (2-oxoglutarate homoexchange) | 2.93 | <u>Smith & Walker, 2003</u> |
| Purified from rat brain mitochondria | 0.64 | <u>De Palma et al., 2010</u> |
| **Recombinant produced in *E. coli*** | **47.24** | **This work** |

